# How mutations of intrinsically disordered protein regions can drive cancer

**DOI:** 10.1101/2020.04.29.069245

**Authors:** Bálint Mészáros, Borbála Hajdu-Soltész, András Zeke, Zsuzsanna Dosztányi

**Affiliations:** MTA-ELTE Momentum Bioinformatics Research Group, Department of Biochemistry, Eötvös Loránd University, Pázmány Péter stny 1/c, Budapest, H-1117 Hungary; Institute of Enzymology, RCNS, Budapest PO Box 7, H-1518, Hungary; EMBL Heidelberg, Meyerhofstraße 1, 69117 Heidelberg, Germany

## Abstract

Many proteins contain intrinsically disordered regions (IDRs) which carry out important functions without relying on a single well-defined conformation. IDRs are increasingly recognized as critical elements of regulatory networks and have been also associated with cancer. However, it is unknown whether mutations targeting IDRs represent a distinct class of driver events associated with specific molecular and system-level properties, cancer types and treatment options. Here, we used an integrative computational approach to explore the direct role of intrinsically disordered proteins/protein regions (IDPs/IDRs) driving cancer. We showed that around 20% of cancer drivers are primarily targeted through a disordered region. The detailed analysis of these IDRs revealed that they can function in multiple ways that are distinct from the functional mechanisms of ordered drivers. Disordered drivers play a central role in context-dependent interaction networks and are enriched in specific biological processes such as transcription, gene expression regulation and protein degradation. Furthermore, their modulation represents an alternative mechanism for the emergence of all known cancer hallmarks independently of the modulation of globular proteins. Disordered drivers are also highly relevant at the sample level, and their mutations can represent the key driving event in certain individual cancer patients. However, treatment options for such patients are currently severely limited. The presented study highlights a largely overlooked class of cancer drivers associated with specific cancer types that need novel therapeutic options.

## Introduction

The identification of cancer driver genes and understanding their mechanisms of action is necessary for developing efficient therapeutics. Many cancer-associated genes encode proteins that are modular, containing not only globular domains but also intrinsically disordered proteins/regions (IDPs/IDRs). IDRs can be characterized by conformational ensembles, however, the detailed properties of these ensembles can vary greatly from largely random/like behavior to exhibiting strong structural preferences, with the length of these segments ranging from a few residues to domain-sized segments. The function of IDRs relies on their inherent conformational heterogeneity and plasticity, enabling them to act as flexible linkers or entropic chains, mediate transient interactions through linear motifs, direct the assembly of macromolecular assemblies or even drive the formation of membraneless organelles through liquid-liquid phase separation[1,2,3,4]. In general, disordered regions are core components of interaction networks and fulfill critical roles in regulation and signaling[5]. In accord with their crucial functions, IDPs are often associated with various diseases[6], in particular with cancer. The prevalence of protein disorder among cancer-associated proteins was observed in general[7]. However, cancer-associated missense mutations showed a strong preference for ordered regions which indicates that the association between protein disorder and cancer might be indirect[8]. Nevertheless, a direct link between protein disorder and cancer was suggested in the case of two common forms of generic alterations; chromosomal rearrangements[9] and copy number variations[10]. Cancer mutations were shown to occur within linear motif sites located in IDRs [11]. In a specific case, the creation of IDR-mediated interactions was suggested to lead to tumorigenesis[12]. However, it has not been systematically analyzed whether mutations of IDRs can have a direct role driving cancer development and what are the main molecular functions and biological processes altered by such events.

In recent years, thousands of human cancer genomes have become available through large-scale sequencing efforts. The collected genetic variations revealed that cancer samples are heterogeneous and contain a large number of randomly occurring, so-called passenger mutations. Therefore, one of the main challenges for the interpretation of cancer genomics data is the identification of driver genes, whose mutations actively contribute to cancer development. When samples are analyzed in combination, various patterns start to emerge that enable the identification of cancer driving genes[13]. These signals can highlight genes which are frequently mutated in specific types of cancer[14,15], biological processes/pathways that are commonly altered in tumor development[16,17] or traits that govern tumorigenic transformation of cells[18]. The positional accumulation of mutations within specific ordered structures, domains, or interactions surfaces was also shown to be strong indicators of cancer driver roles[19–23]. The number of driver genes is currently estimated to be in the low to mid-hundreds[24], but this number could increase with the growing number of sequenced cancer genomes[15]. However, most of the known, well-characterized driver genes are associated with ordered domains of proteins.

The complex relationship between protein disorder and cancer can be demonstrated through two well-characterized examples, p53 and β-catenin. As a tumor suppressor, p53 is most commonly altered by truncating mutations, but it also contains a large number of missense variations. Mutations collected from multiple patients across different cancer types tend to cluster within the central region which corresponds to the ordered DNA-binding domain of p53 [25]. In contrast, significantly fewer mutations correspond to the disordered N- and C-terminal regions - which are involved in numerous, sometimes overlapping protein-protein interactions [26]. In particular, almost no mutations are located within the N-terminal region corresponding to a so-called degron motif, a linear motif site recognized by the E3 ligase MDM2 that plays a critical role in regulating the degradation of p53 [27]. Furthermore, the tetramerization domain in the C-terminal part is also less affected by cancer mutations. This region represents a so-called disordered domain, a conserved region that forms a well-defined structure in its oligomeric form. The tetrameric ordered structure masks a nuclear export signal, which needs to become exposed for the proper function of p53, highlighting the intrinsic dynamical properties of this region [28]. The oncogenic β-catenin presents a completely different scenario. In terms of domain organization, β-catenin also contains a disordered N- and C-terminal and an ordered domain in between [29]. However, in this case the cancer mutations are largely localized to a short segment within the N-terminal disordered region which corresponds to the key degron motif regulating the cellular level of β-catenin in the absence of Wnt signalling [8,11].

The aim of this work was to explore if other IDRs, similarly to β-catenin, play a potential driver role in cancer. Based on cancer mutations collected from genome-wide screens and targeted studies[30], we identified significantly mutated protein regions[31] and classified them into ordered and disordered regions integrating experimental structural knowledge and predictions. Automated and high-quality manually curated information were gathered for the collected examples to gain better insights into their functional and system-level properties, and confirm their roles in tumorigenic processes. We aimed to answer the following questions: What are the characteristic molecular mechanisms, biological processes, and protein-protein interaction network roles associated with proteins mutated at IDRs? And at a more generic level: how fundamental is the contribution of IDPs to tumorigenesis? Are IDP mutations just accessory events, or can they be the dominant molecular background to the emergence of cancer? Is there a characteristic difference in terms of treatment options between patient samples targeted mostly within ordered and disordered regions?

## Results

### Disordered protein modules are targets for tumorigenic mutations

For the purpose of our analysis it was necessary to use an approach that can identify not only cancer drivers, but also the specific regions directly targeted by cancer mutations. We used the iSiMPRe[31] method, which can highlight significantly mutated regions without prior assumptions about the type or the size of such regions and was shown to perform similarly to other methods in identifying cancer drivers [32]. Cancer mutations were collected from the COSMIC and TCGA databases and were pre-filtered to minimize the potential random accumulation of mutations that IDRs are more prone to [31]. We restricted our analysis to high-confidence cases to minimize the chance of false positives. The order/disorder status of the identified significantly mutated regions was determined based on experimental data or homology transfer, when available, or by using a combination of prediction approaches (See Data and Methods). Cancer drivers were manually characterized as tumor suppressors, oncogenes and context-dependent cancer genes based on the literature.

Altogether, we identified 178 ordered and 47 disordered driver regions in 145 proteins from the human proteome (Supplementary Table S1, Figure 1A). The ratio of disordered driver regions is lower than expected on the ratio of disordered residues (21% vs 30%). This is the case for both oncogenes and tumor suppressor genes, but not for context-dependent genes. Further underlining the relevance of IDRs, context-dependent cancer drivers also have more residues and mutations within disordered regions in general together with a slightly higher proportion of disordered drivers (see Supplementary Figure S1).

**Figure 1.**
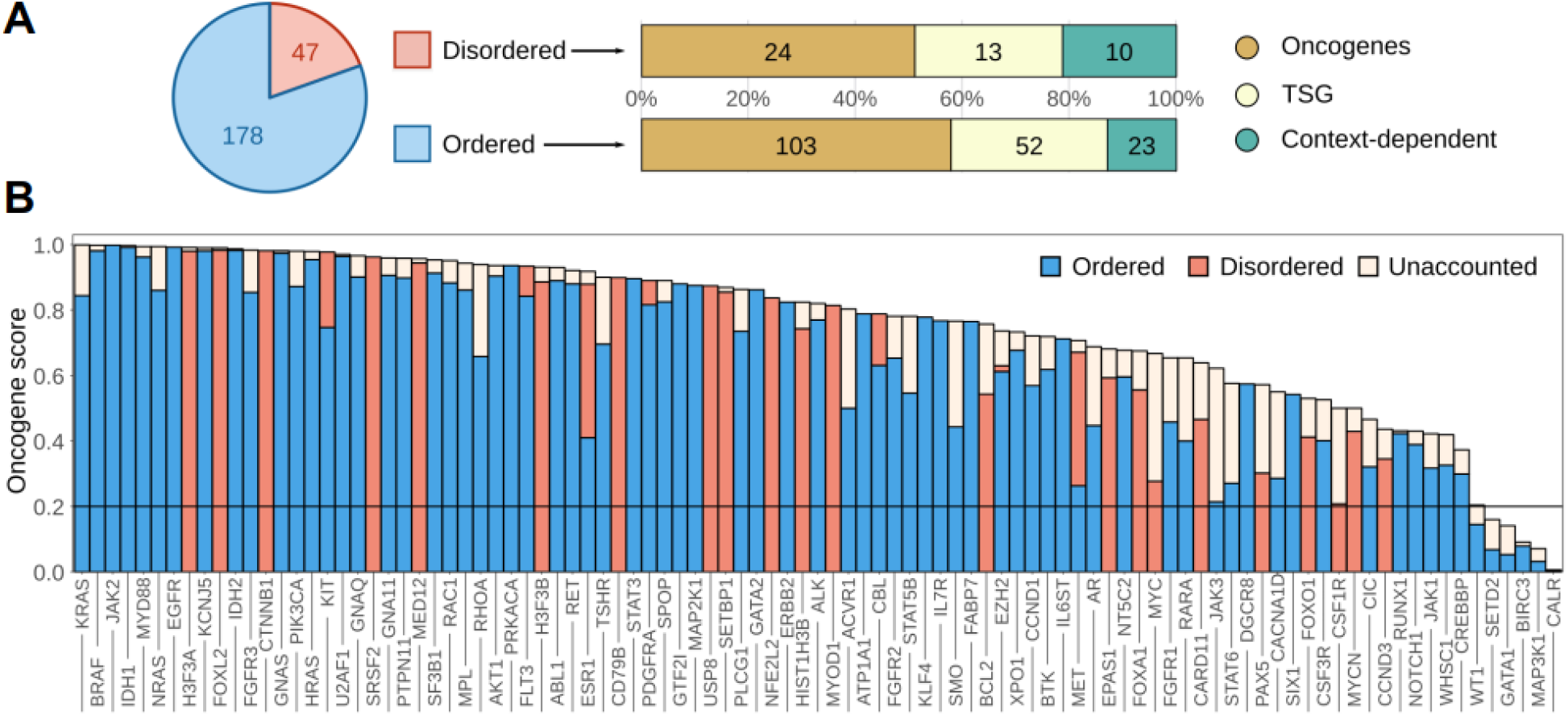
The distribution of ordered and disordered driver protein regions. A: The distribution of ordered and disordered driver protein regions. B: Oncogene scores of full genes and oncogene scores explained by the identified regions in oncogenes and context-dependent driver genes. ‘Unaccounted’ corresponds to the fraction of mutations not in the identified, high significance regions.

The identified driver regions typically represent compact modules, usually not covering more than 10% or 20% of the sequences in the case of oncogenes and tumor suppressors, respectively (Supplementary Figure S2). It was suggested that true oncogenes are recognizable from mutation patterns according to the 20/20 rule, having a higher than 20% fraction of missense point mutations in recurring positions (termed the oncogene score[33]). In contrast, tumor suppressors have lower oncogene scores, and predominantly contain truncating mutations. Figure 1B shows that the 20/20 rule holds true for the vast majority of the identified region-harboring oncogenes and context-dependent genes, even based on the oncogene scores calculated from the identified regions alone. This underlines that the identified driver regions are the main source of the oncogenic effect in almost all cases. While most drivers contain both ordered and disordered modules, oncogenesis is typically mediated through either ordered or disordered mutated regions. This effectively partitions cancer drivers into ‘ordered drivers’ and ‘disordered drivers’, regardless of the exact structural composition of the full protein.

While many of the disordered drivers have already been identified previously as cancer drivers, our analysis identified 13 additional examples that were not included in the list identified cancer drivers collected recently [24]. However, even in these cases there is literature data supporting the importance in driving cancer (Supplementary Table S2).

### Disordered drivers function via distinct molecular mechanisms

We collected available information about the possible mechanisms of action altered in cancer (Figure 2, Supplementary Table S2). Although this information was partially incomplete in several cases, it still allowed us to highlight the distinct properties of the identified disordered drivers.

**Figure 2.**
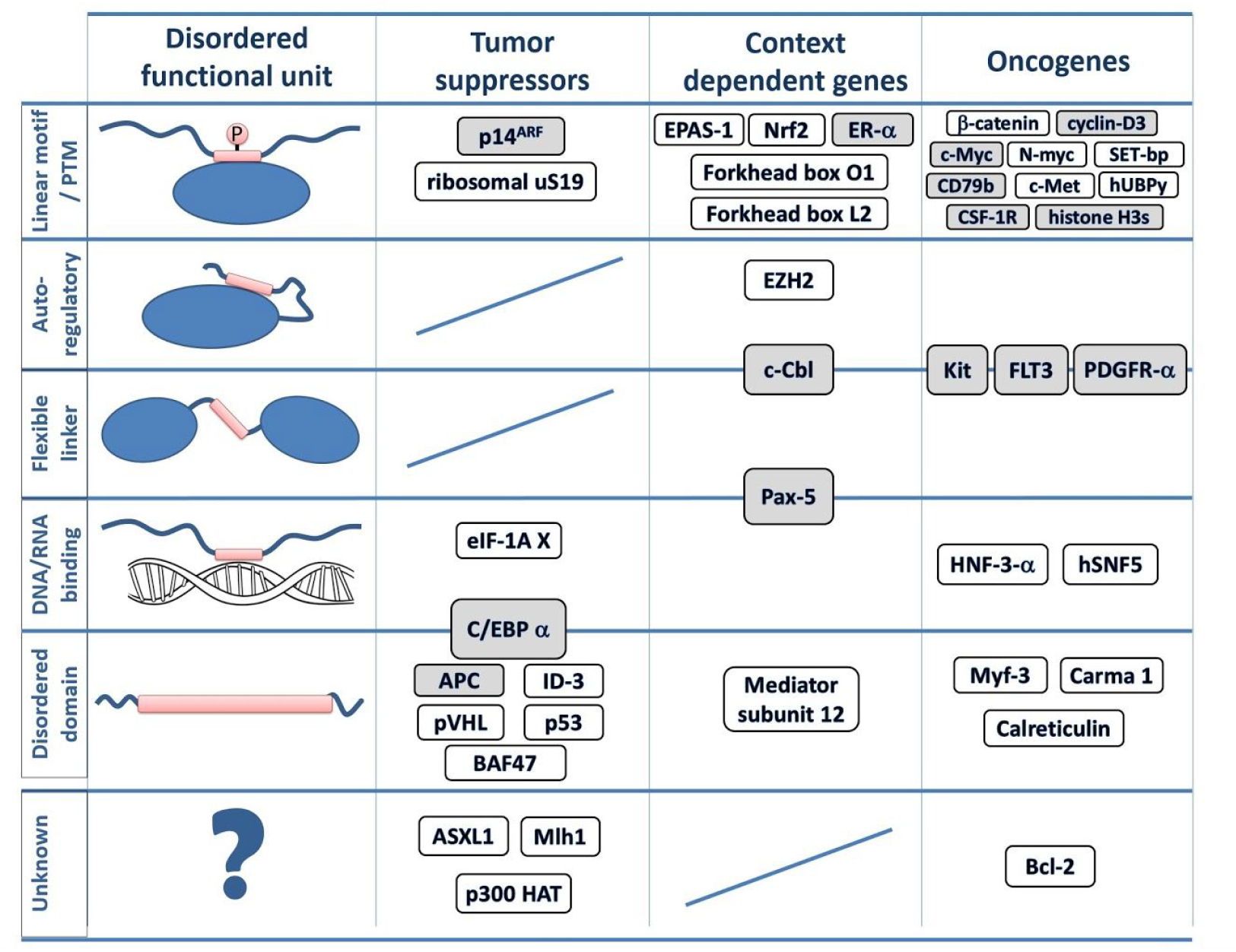
IDP regions mutated in cancer. The classification of identified disordered cancer drivers. Protein names in gray indicate known switching mechanisms either via PTMs or overlapping functions. In protein architecture schematics, blue ovals represent folded domains, blue lines disordered regions and red rectangles represent disordered driver modules. For detailed mutation profiles for each gene, see online visualization links in Supplementary Table S2.

Several of the identified highly mutated disordered regions correspond to **linear motifs**, including sites for protein-protein interactions (e.g. hUBPy [corresponding gene: USP8], forkhead box protein O1 [FOXO1] and ER-α [ESR1]) or degron motifs that regulate the degradation of the protein (e.g β-catenin [CTNNB1], cyclin-D3 [CCND3] and CSF-1R [CSF1R]). However, other types of disordered functional modules can also be targeted by cancer mutations. **IDRs with autoinhibitory roles** (e.g. modulating the function of adjacent folded domains) are represented by EZH2 [EZH2], a component of the polycomb repressive complex 2. While the primary mutation site in this case is located in the folded SET domain, cancer mutations are also enriched within the disordered C-terminus that normally regulates the substrate binding site on the catalytic domain. Another category corresponds to **regions involved in DNA and RNA binding**. The highly flexible C-terminal segment of the winged helix domain is altered in the case of HNF-3-α [FOXA1], interfering with the high affinity DNA binding. For the splicing factor hSNF5 [SRSF2], mutations affect the RNA binding region (Figure 2).

Larger functional disordered modules, often referred to as **intrinsically disordered domains** (IDDs), can also be the primary sites of cancer mutations. Mutated IDDs exhibit varied structure and sequence features. In pVHL [VHL], the commonly mutated central region adopts a molten globule state in isolation[34]. The mutated region of APC [APC] incorporates several repeats containing multiple linear motif sites, which are likely to function collectively as part of the β-catenin destruction complex[35]. In calreticulin [CALR], cancer mutations alter the C-terminal domain-sized low complexity region, altering Ca^2+^ binding and protein localization[36].

**Linker IDRs**, not directly involved in molecular interactions, are also frequent targets of cancer mutations. The juxtamembrane regions located between the transmembrane segment and the kinase domain of Kit [KIT] and related kinases, are the main representatives of this category. Similarly, the regulatory linker region connecting the substrate- and the E2 binding domains is one of the dominant sites of mutations in the case of the E3 ubiquitin ligase c-Cbl [CBL].

One of the recurring themes among cancer-related IDP regions is the formation of molecular switches (Supplementary Table S2). The most commonly occurring switching mechanism involves various post-translational modifications (PTMs), including serine or threonine phosphorylation (e.g. cyclin-D3 [CCND3], c-Myc [MYC] and APC [APC]), tyrosine phosphorylation (e.g. c-Cbl [CBL], CD79b [CD79B], and CSF-1R [CSF1R]), methylation (e.g. histone H3s [H3F3A/H3F3B/HIST1H3B]) or acetylation (e.g. ER-α [ESR1]). An additional way of forming molecular switches involves overlapping functional modules (Figure 2). In the case of BAF47 [SMARCB1], the mutated inhibitory sequence is likely to normally also mask a nuclear export signal in the autoinhibited state[37]. In the case of Pax-5 [PAX5], the mutated flexible linker region is also involved in the high affinity binding of the specific DNA binding site[38]. Cancer mutations of the bZip domain of C/EBP-α [CEBPA] disrupt not only the DNA binding function, but the dimerization domain as well[39]. In addition to their linker function, the juxtamembrane regions of kinases are also involved in autoinhibition and trans-phosphorylation, regulating degradation and downstream signaling events[40,41].

The collected examples of disordered regions mutated in cancer cover both oncogenes and tumor suppressors, as well as context-dependent genes (Figure 2). There is a slight tendency for tumor suppressors to be altered via longer functional modules, such as IDDs. Nevertheless, with the exception of linkers in tumor suppressors and IDDs in context-dependent genes, every other combination occurs even within our limited set.

### Disordered driver mutations preferentially modulate receptor tyrosine kinases, DNA-processing and the degradation machinery

Disordered and ordered drivers can employ different molecular mechanisms in order to fulfill their associated biological processes. To quantify these differences, we assembled a set of molecular toolkits integrating Gene Ontology terms (see Data and methods and Supplementary Table S3). Based on this, we calculated the enrichment of each molecular toolkit in both disordered and ordered drivers, in comparison with the full human proteome, highlighting enriched and possibly driver class-specific toolkits (Figure 3A). Receptor activity is the most enriched function for both types of drivers, owing at least partially to the fact that receptor tyrosine kinases can often be modulated via both ordered domains and IDRs (Figure 1B). In contrast, the next three toolkits enriched for disordered drivers are highly characteristic of them. These are gene expression regulation and the modulation of DNA structural organization - together representing the structural and the information content-related aspects of DNA processing -, and the degradation of proteins, mainly through the ubiquitin-proteasome system. In addition, RNA processing, translation and folding is also characteristic of disordered drivers; and while this toolkit is not highly enriched compared to the human proteome in general, ordered drivers are almost completely devoid of this toolkit.

**Figure 3.**
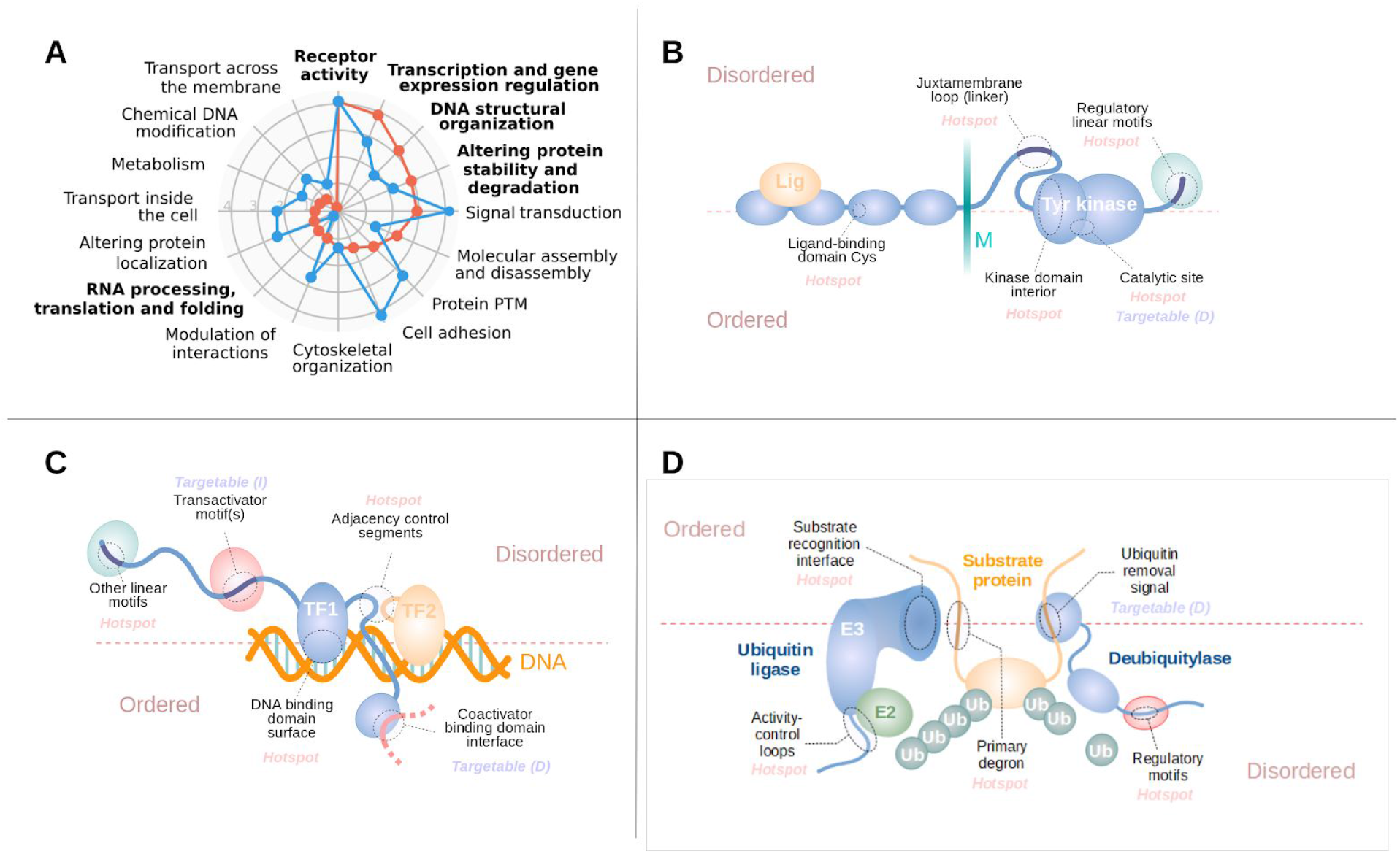
Pathways and processes modulated by disordered driver mutations. A: Overrepresentation of molecular toolkits defined based on GO terms for ordered (blue) and disordered (red) drivers, compared to the average of the whole human proteome. B-D: schematic examples of RTKs (B), transcription factors (C) and components of the ubiquitin ligase machinery (D) that are modulated through disordered driver regions.

Among the highlighted functional groups, receptor tyrosine kinases (RTKs) are well-known to be major players in tumorigenesis[42]. While for several RTKs, the major mutational events are oncogenic kinase domain mutations, there are also RTKs that contain a secondary disordered mutation site with lower incidence rates, or an alternative primary site, where the main target for mutations is context-dependent. Several RTKs are clear examples of this context dependence: gastrointestinal stromal tumor mutations prefer IDR mutations in both Kit and PDGFR-α[43], while leukaemia prefers catalytic site mutations in Kit. Group III receptor tyrosine kinases in general (including Kit, FLT3 and PDGFR-α) are especially prone to be mutated at their disordered juxtamembrane regions (Figure 3B). In some cases, such as FLT3, these IDRs are the main sites for tumorigenic mutations[44]. However, RTK IDR mutations are not restricted to group III receptor tyrosine kinases, as c-Met also often harbors missense mutations at its juxtamembrane loop region as well. These mutations include missense mutations affecting the Tyr1010 phosphorylation site and exon 19 skipping, removing a degron located within this region[45]. In contrast, for CSF-1R mutations accumulate in the negative regulatory motifs (a c-Cbl ubiquitin ligase binding motif) in the receptor tail, leading to the overactivation of the receptor[46] in various haematopoietic cancers.

Cancer mutations often target various elements of the transcriptional machinery, including transcription factors, repressors, transcriptional regulators, and coactivators/corepressors[47](Figure 3C). In most cases transcription factors are targeted through linear motifs that regulate the degradation (EPAS-1, β-catenin, c-Myc and N-myc) or localization of the protein (Forkhead box protein O1). Mutated IDRs can also directly affect the activity of the protein. These regions often work in conjunction with a separate DNA-binding domain, and can shift affinities for various DNA-binding events (such as HNF-3-α mutations preferentially affecting low-affinity DNA binding[48]), or can disrupt interaction with cofactors (such as the SMAD3 interaction of the Forkhead box protein L2[49]). In the case of bZip-type dimeric transcription factors, mutations can affect the interaction through the modulation of the disordered dimerization domain. Depending on the activating/repressive function of individual transcription factors, IDR-mutated proteins can be both oncogenes (with Myf-3 mutations promoting the dimerization with c-Myc[50]), or tumor suppressors (with ID-3 mutations impairing its repressor activity[51]). Disordered mutational hotspots also target other elements of the transcription machinery, affecting either covalent or non-covalent histone modifications, altering histone PTMs or histone exchange/movement along the DNA. However, the exact role of several other proteins involved in chromatin remodelling is still somewhat unclear (SETBP1, or ASXL1).

The alteration of protein abundance through the ubiquitin-proteasomal system (UPS) is a central theme in tumorigenesis [27]. Interestingly, ubiquitination sites are seldom mutated directly, while in several cases, cancer mutations directly alter degron motifs which typically reside in disordered protein regions (Figure 3D). Such mutations lead to increased abundance of the protein by disrupting the recognition by the corresponding E3 ligase. Complementing degron mutations, ubiquitin ligases are also implicated in tumorigenesis (Figure 3D). These enzymes are typically highly modular and can harbor driver mutations in both ordered and disordered regions (Supplementary Table S1). FBXW7 is mutated at its ordered substrate-binding domain, paralleled with target degron mutations in its substrates, c-Myc and N-myc. In contrast, pVHL, which is the substrate recognition component of the cullin-2 E3 ligase complex, is targeted through a large disordered driver region, with its target EPAS-1 bearing a mutant degron. The activity of c-Cbl, the main E3 ligase responsible for the regulation of turnover for RTKs, is targeted through a disordered linker/autoregulatory region in AML and other hematopoietic disorders. In addition to the disruption of ubiquitination, the enhancement of deubiquitination can also provide a tumorigenic effect. hUBPy, the deubiquitinase required for entry into the S phase, is mutated at its disordered 14-3-3-binding motif, enhancing deubiquitinase activity in lung cancer[52].

### Disordered mutations give rise to cancer hallmarks by targeting central elements of biological networks

Almost all of the analyzed IDRs are involved in binding to a molecular partner, even some of the linkers owing to their multifunctionality. Therefore, we analyzed known protein-protein interactions of ordered and disordered cancer drivers in more detail (see Data and methods). Our results indicate that both sets of drivers are involved in a large number of interactions, and show increased betweenness values compared to average values of the human proteome, and even compared to the direct interaction partners of cancer drivers (Figure 4A). However, this trend is even more pronounced for disordered drivers. The elevated interaction capacity could also be detected at the level of molecular function annotations using Gene Ontology (see Supplementary Table S4 and Data and methods). Figure 4B shows the average number of types of molecular interaction partners for both disordered and ordered drivers, contrasted to the average of the human proteome. The main interaction partners are similar for both types of drivers, often binding to nucleic acids, homodimerizing, or binding to receptors. However, disordered drivers are able to physically interact with a wider range of molecular partners, and are also able to more efficiently interact with RNA and the effector enzymes of the post-translational modification machinery. This, in particular, can offer a way to more easily integrate and propagate signals through the cell, relying on the spatio-temporal regulation of interactions via previously demonstrated switching mechanisms (Supplementary Table S2).

**Figure 4.**
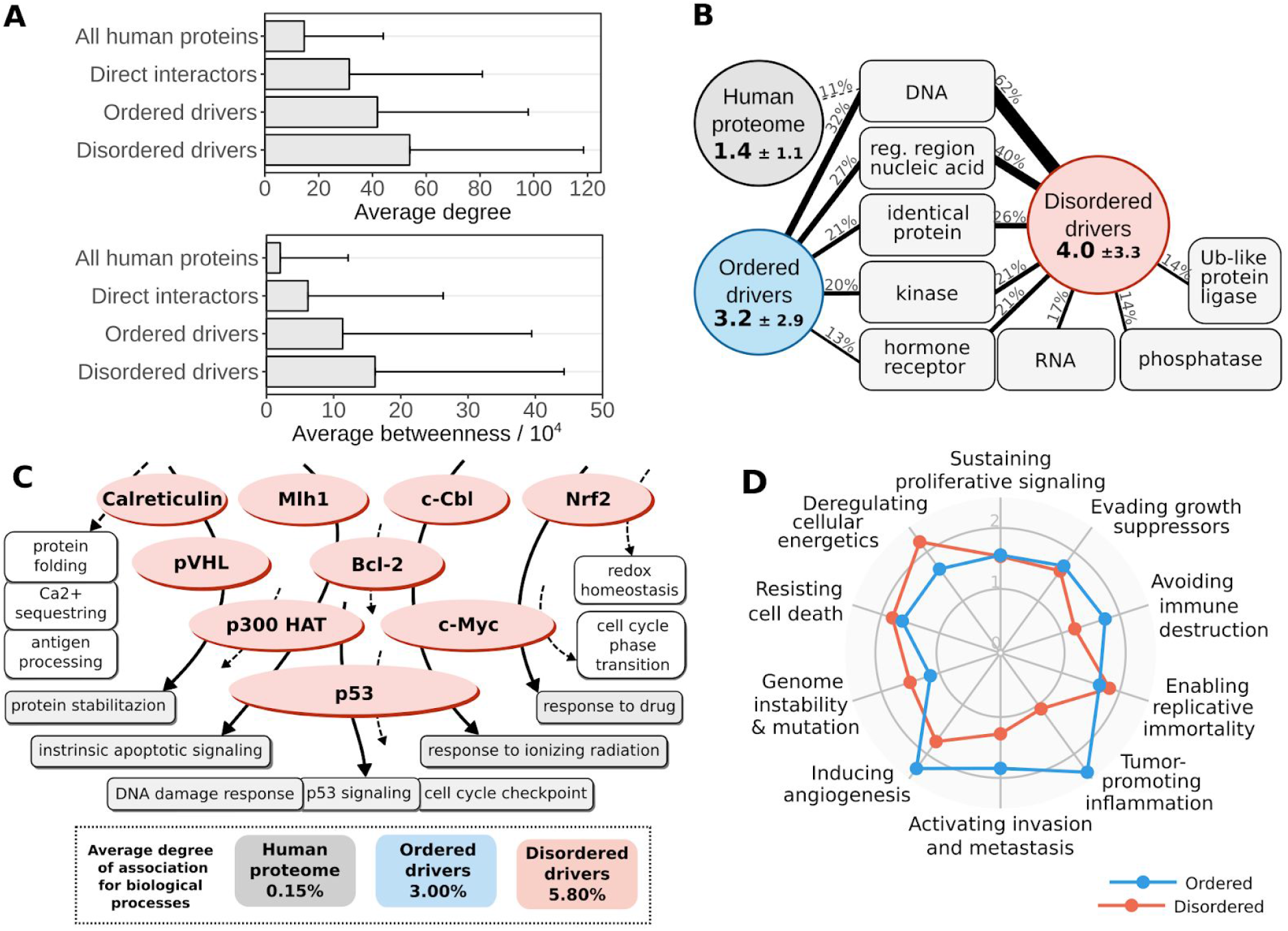
Characteristics of cancer drivers at the network/pathway-, and cellular levels. A: Average degree (top) and betweenness (bottom) of all human proteins, the direct interaction partners of drivers, ordered drivers, and disordered drivers. B: the average occurrence of various types of interaction partners for the whole human proteome (grey circle), ordered drivers (blue circle) and disordered drivers (red circle). Values in circles show the average number of types of interactions together with standard deviations. The most common interaction types are shown in grey boxes with connecting lines showing the fraction of proteins involved in that binding mode. Only interaction types present for at least 1/8th of proteins are shown. C right: average values of overlap between protein sets of various biological processes, considering the full human proteome (grey), ordered drivers (blue) and disordered drivers (red). Left: an example subset of disordered drivers with associated biological processes marked with arrows (dashed and solid arrows marking processes involving only one, or several disordered drivers). Process names in grey represent processes that involve at least two disordered drivers, names in white boxes mark other processes attached to disordered drivers. D: Overrepresentation of hallmarks of cancer for ordered (blue) and disordered (red) drivers compared to all census drivers.

The high interaction capacity and central position of disordered drivers allows them to participate in several biological processes. The association between any two processes can be assessed by quantifying the overlap between their respective protein sets (see Data and methods). We analyzed the average overlap between various processes using a set of non-redundant human-related terms of the Gene Ontology (Supplementary Table S5). The average overlap of proteins for two randomly chosen processes is 0.15%, showing that as expected, in general, biological processes utilize characteristically different gene/protein sets. Restricting proteins to the identified drivers, and only considering processes connected to at least one of them, the average overlap between processes is increased to 3.00% for ordered drivers and 5.80% for disordered drivers (Figure 4C). This shows that the integration of various biological processes is a distinguishing feature of cancer genes in general, and for disordered drivers in particular; and that IDPs targeted in cancer are efficient integrators of a wide range of processes.

Cancer hallmarks describe ubiquitously displayed traits of cancer cells[18]. In order to quantify the contribution of drivers to each of the ten hallmarks, we manually curated sets of biological process terms from the Gene Ontology that represent separate hallmarks (see Data and methods and Supplementary Table S6). Enrichment analysis of these terms shows that all hallmarks are significantly over-represented in census cancer drivers compared to the human proteome (Supplementary Figure S3A), serving as a proof-of-concept for the used hallmark quantification scheme. Furthermore, comparing drivers with identified regions to all census cancer drivers shows a further enrichment (Supplementary Figure S3B), indicating that the applied region identification protocol of iSiMPRe is able to pick up on the main tumorigenic signal by pinpointing strong driver genes. Separate enrichment calculations for ordered and disordered drivers show that despite subtle differences in enrichments, in general, all ten hallmarks are over-represented in both driver groups (Figure 4D). This indicates that while the exact molecular mechanisms through which ordered domain and IDR mutations contribute to cancer are highly variable, both types of genetic modulation can give rise to all necessary cellular features of tumorigenic transformation. Hence, IDR mutations provide a mechanism that is sufficient on its own for cancer formation.

### Disordered drivers can be the dominant players at the patient sample level

We assessed the role of the identified drivers at the patient level using whole-genome sequencing data from TCGA 10,197 tumor samples containing over three and a half million genetic variations were considered to delineate the importance of disordered drivers at the sample level across the 33 cancer types covered in TCGA. In driver region identification we only considered mutations with a local effect (missense mutations and in frame indels), which naturally yielded only a restricted subset of all true drivers. However, in patient level analyses, we also considered other types of genetic alterations of the same gene, in order to get a more complete assessment of the alteration of identified driver regions per cancer type (see Data and methods).

In spite of the incompleteness of the identified set of driver genes, we still found that on average about 80% of samples contain genetic alterations that affect at least one identified ordered or disordered driver region. Thus, the identified regions are able to describe the main players of tumorigenesis at the molecular level (Figure 5A). While at the protein level typically either ordered or disordered regions are modulated (Figure 1B), at the patient level most samples show a mixed structural background, most notably in colorectal cancers (COAD and READ). Some cancer types, however, show distinct preferences for the modulation of a single type of structural element: for thyroid carcinoma (THCA) or thymoma (THYM) the molecular basis is almost always the exclusive mutation of ordered protein regions. At the other extreme, the modulation of disordered regions is enough for tumor formation in a considerable fraction of cases of liver hepatocellular, adrenocortical, and renal cell carcinomas, together with diffuse large B-cell lymphoma (LIHC, ACC, KIRC and DLBC). These results, in line with the previous hallmark analyses, show that IDR mutations can constitute a complete set of tumorigenic alterations. Hence, there are specific subsets of patients that carry predominantly, or exclusively disordered driver mutations in their exome.

**Figure 5:**
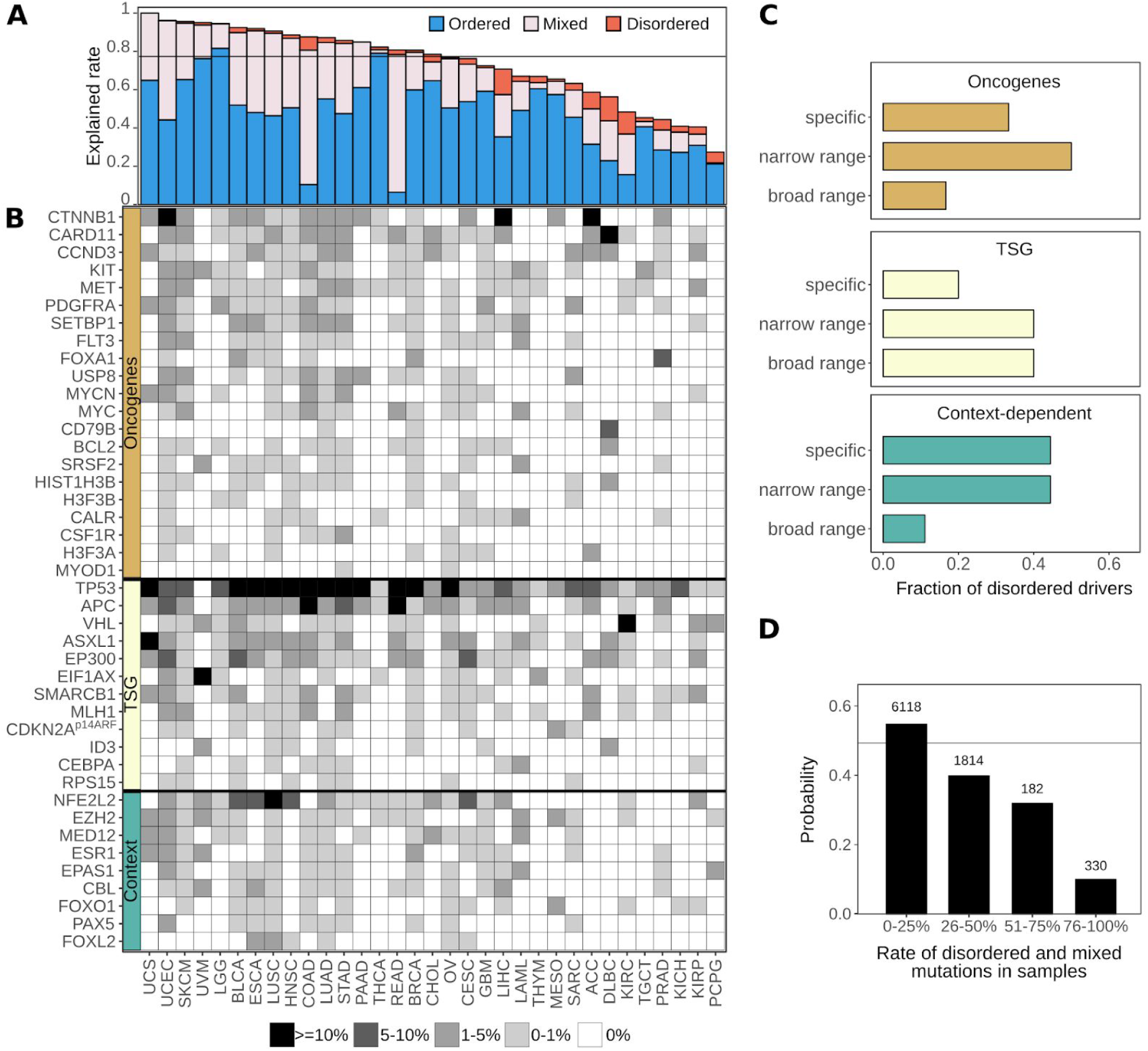
Therapeutic options for targeting disordered drivers. A: Fraction of samples that contain altered driver genes per cancer types. Samples can contain mutations affecting only ordered drivers (blue), only disordered drivers (red) or both (mixed - gray). B: Percentage of cancer samples, grouped by cancer types, containing genetic alterations that target the identified disordered driver regions. C: The distribution of disordered drivers from the three classes of cancer genes categorized into specific, narrow and broad range, based on the frequency of samples they are mutated in (see Data and methods). D: The probability of having an available FDA approved drug for at least one mutation-affected gene for patients, as a function of the ratio of affected disordered genes compared to all mutated genes in the sample. The horizontal black line represents the total fraction of targetable samples (0.49) from 8,444 samples.

Whole genome sequencing data was also used to assess the cancer type specificity of disordered drivers (Figure 5B). Basically, all studied cancer types have at least one disordered driver that is mutated in at least 1% of cases, with the exception of thyroid carcinoma (THCA). There are only four disordered drivers that can be considered as generic drivers, being mutated in a high number of cancer types. p53 presents a special case in this regard, as it is the main tumor suppressor gene in humans, and thus is most often affected by gene loss or truncations which are likely to eliminate the corresponding protein product. These alterations abolish the function of both the ordered and disordered driver regions at the same time (the DNA-binding domain and the tetramerization region). In contrast, the other three generic disordered drivers are predominantly altered via localized mutations in their disordered regions: the degrons of β-catenin and NRF2, and the central region of APC, and hence these are true disordered drivers which are commonly mutated in several cancer types. However, the majority of disordered drivers show a high degree of selectivity for tumor types, being mutated only in very specific cancer types. This specificity is strongly connected to the tumorigenic roles of disordered drivers (Figure 5C). Considering 1% of patient samples as the cutoff, tumor suppressors are typically implicated in a broad range of cancer types, while oncogenes on average show a high cancer type specificity. Context-dependent disordered drivers are often mutated in only a very restricted set of cancers.

Strikingly, the identified disordered drivers can have an even more dominant role. In several rarer cancers or more specific cancer subtypes which are not included in the broad classes described in TCGA (including both malignant and benign cases), mutations in a specific disordered driver is the main, or one of the main driver events (Table 1). Altogether, this list involves 18 of our disordered cancer drivers. In the collected cancer types, targeting disordered regions can have a potentially huge treatment advantage, and in many cases, the counteraction of these IDR mutations may be the only viable therapeutic strategy.

**Table 1:**
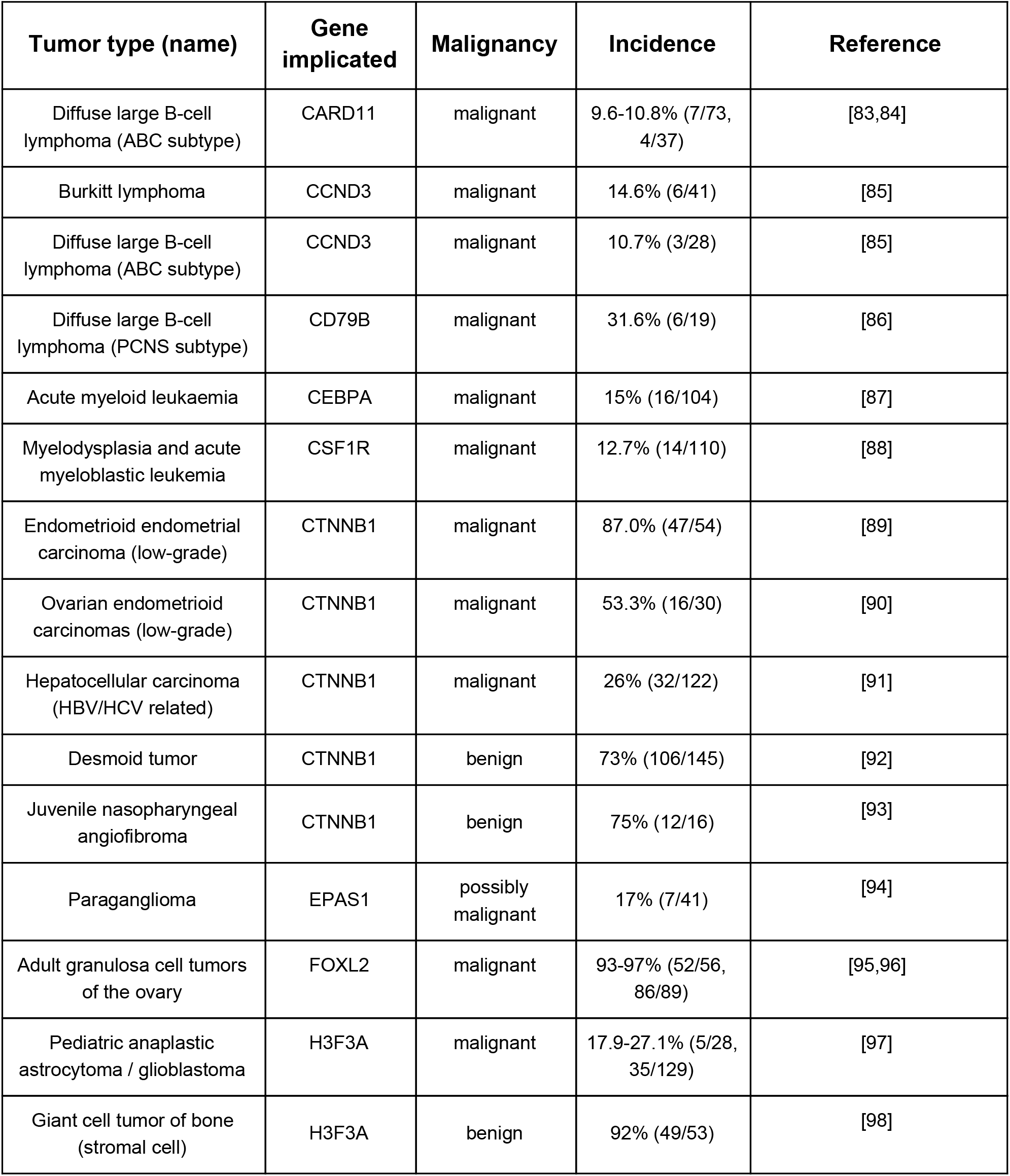
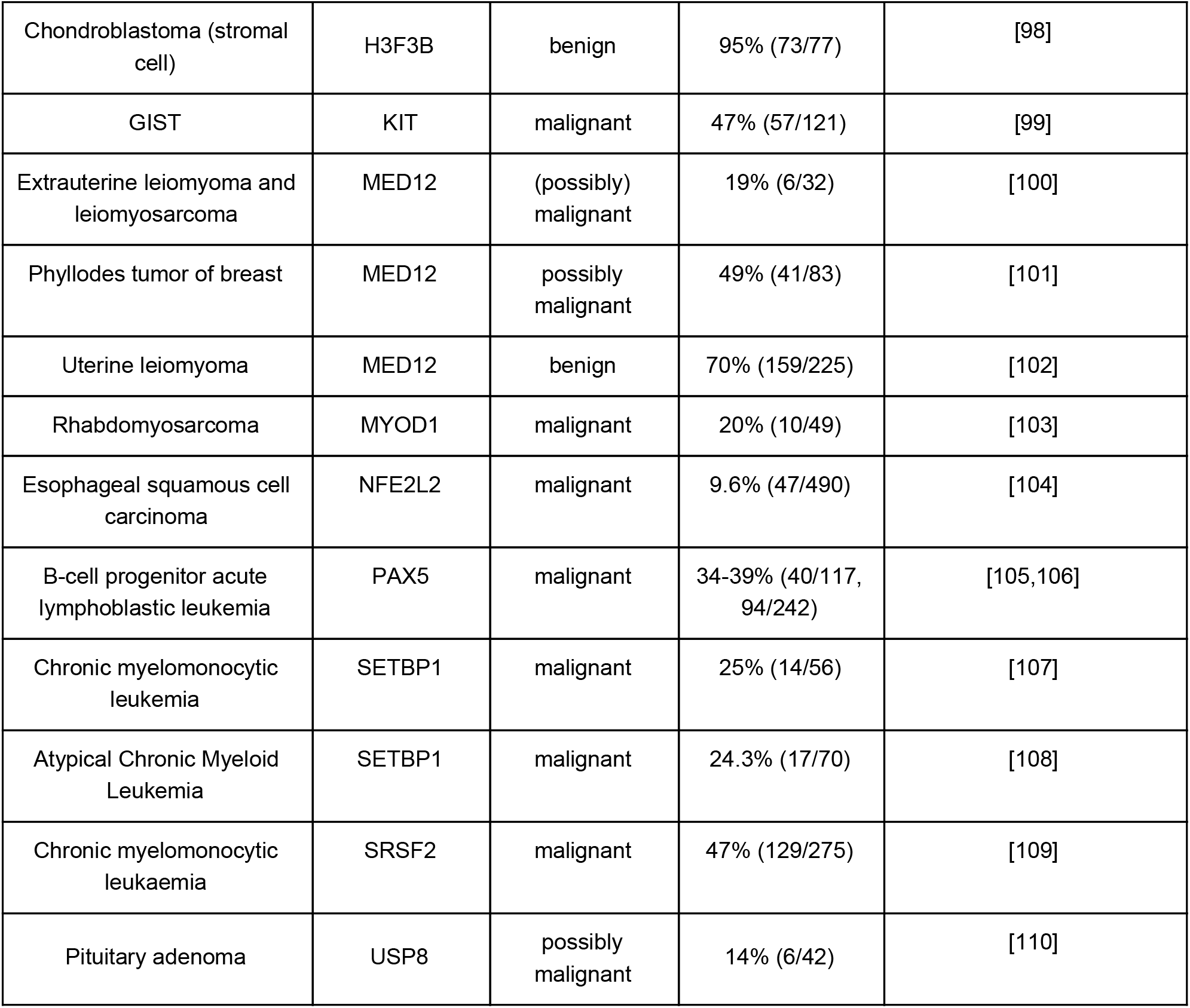
Cancer types with mutation incidence rates around or above 10% in the disordered driver gene of interest per total patients studied.

### Cancer incidences arising through disordered drivers lack effective drugs

Next, we addressed how well disordered drivers are targetable by current FDA approved drugs, as collected by the OncoKB database[53]. This database currently contains 83 FDA-approved anticancer drugs, either as part of standard care or efficient off-label use (see Data and methods). These drugs have defined exome mutations that serve as indications for their use. The majority of these drugs target ordered domains, mostly inhibiting kinases. Currently only 7 drugs are connected to disordered region mutations, which correspond to only four sites in FGFR and c-Met. These drugs act indirectly, targeting ordered kinase domains, to counteract the effect of the listed activating disordered mutations.

This represents a clear negative treatment option bias against patients whose tumor genomes contain disordered drivers. Considering all mutations in patient samples gathered in TCGA, the fraction of disordered driver mutations actually serves as an indicator of whether there are suitable drugs available. Patients with mostly ordered driver mutations have a roughly 50% chance that an FDA-approved drug can be administered with expected therapeutic effect. This chance drops to 10% for patients with predominantly disordered mutations (Figure 5D). Thus, incidences of cancer arising through disordered driver mutations are currently heavily under-targeted, highlighting the need for efficient targeting strategies for IDP driven cancers.

## Discussion

In recent years, cancer genome projects revealed the genomic landscapes of many common forms of human cancer. As a result, several hundred cancer driver genes have been identified whose genetic alterations can be directly linked to tumorigenesis[24]. Only a few of these genes correspond to “mutation mountains”, ie. genes that are commonly altered in different tumor types, while most of the cancer drivers altered infrequently [33]. Cancer driver genes are associated with a set of core cellular processes, also termed hallmarks [18]. At a more detailed level, however, drivers are surprisingly heterogeneous in terms of molecular functions and cellular roles. In this work we showed that cancer drivers are also diverse in terms of their structural properties. Using an integrated computational approach we identified a set of cancer drivers that are specifically targeted by mutation in a disordered region. IDRs represent around 30% of residues in the human proteome and are also an integral part of many cancer-associated proteins. Despite the critical roles of these regions, they are often not the main sites of driver mutations [8]. Our results confirmed that driver mutations that alter the proper functioning of ordered domains of the encoded protein are slightly overrepresented compared to those that modulate the function of disordered regions. Nevertheless, in a significant number of cases, corresponding to around 20% of the mutated drivers, cancer mutations specifically target disordered regions (Figure 1A).

The critical role of these disordered drivers in tumorigenesis is supported not only by the enrichment of single nucleotide variations and in-frame insertions and deletions, but also by literature data (Supplementary Table 2). Disordered drivers are associated with known cancer hallmarks through specific biological processes (Figure 3A) and show strong evolutionary conservation [54]. Driver mutations within IDRs are present in samples across a wide range of cancer types, and can also be the main, or one of the main driver events for several tumor subclasses, including both malignant and benign cases (Table 1). Our work highlighted several novel drivers that are not yet included in the previous collections of cancer driver genes previously assembled based on a combination of computational methods [24], indicating a hidden bias in the identification of driver genes.

The collection of disordered cancer drivers highlighted many interesting examples that carry out important functions without relying on a well-defined structure, extending the list of IDR with disease relevance. Many of the collected cases correspond to linear motif sites which mediate interactions with globular domains, regulating interactions, localization or cellular fate of proteins. However, the collected examples represent a broader set of functional mechanisms, encompassing DNA and RNA binding regions, linkers, autoinhibitory segments and disordered domains. These functional modules can also regulate the assembly of large macromolecular complexes and regulate the activity of neighboring domains. The key to the proper functioning of the targeted IDRs is their structural disorder which enables them to undergo drastic conformational changes depending on context-dependent regulation. While in most cases it has been characterized how mutations of the critical IDR disrupts the balance between the different functional states, our understanding of this mechanism is still incomplete for several examples (Figure 2, Supplementary Table 2). For instance, the mutation and conservation pattern of MLH1 highlights a novel linear motif site within the disordered linker region of MLH1 with unknown function. In the case of p14Arf, the functional role of the mutated region needs to be revisited in the light of recent evidence on the relevance of phase separation organizing the nucleolus [55]. ASXL1 and p300 HAT are both involved in chromatin remodelling, but little is known about the functional roles of the disordered regions targeted by cancer mutations.

At the patient level, samples in general contain a combination of genetic alterations that involves both ordered and disordered drivers. However, patients with mostly IDR mutations typically have much more limited treatment options. Most current anticancer drugs target ordered protein domains, and are inhibitors designed against enzyme activity (using either competitive or noncompetitive inhibition)[56–58]. In general, currently successful drug development efforts mainly focus on ordered protein domains, derived within the framework of structure-based rational drug design[59]. However, IDPs can potentially offer new directions for cancer therapeutics[60]. Currently tested approaches include the direct targeting of IDPs by specific small compounds, or blocking the globular interaction partner of IDPs[61,62]. The successful identification of disordered drivers and corresponding tumor types provides the first step in providing the means for new therapeutic interventions in cancer types that currently lack treatment options.

In conclusion, in this work, we went beyond a simple association between IDRs and cancer by taking advantage of the avalanche of data produced by systematic analyses and large-scale sequencing projects of cancer genomes. Our work underlines the direct driver role of IDRs in cancer. It provides fundamental insights into the specific molecular mechanisms and regulatory processes altered by cancer mutations targeting IDRs, highlighting important regions that need further structural and functional characterizations. Furthemore, we showed that many already known cancer drivers rely on intrinsic flexibility for their function and identified novel cancer drivers that had been overlooked by current driver identification approaches, revealing a structure-centric bias that still exists in these methods. Importantly, our work also demonstrates the relevance of disordered drivers at the patient level and highlighting a strong need to expand treatment options for IDRs. By looking at the timeline of the COSMIC database, we can observe a steady growth of disordered drivers with every new release (Supplementary Figure S4). Nevertheless, our study was restricted to cases that were targeted by point mutations or in-frame insertions or deletions, therefore, the location of alterations can be directly linked to the perturbed functional module. However, there are additional disordered drivers that are altered via more complex genetic mechanisms in cancer, such as specific frameshift mutations (e.g. NOTCH1[63]), chromosomal translocations (e.g. BCR[64], ERG [65]), or copy number variations (e.g. p14ARF[66]). Altogether, these observations suggest that we can expect the emergence of further examples of genetic alterations of driver genes that interfere with structurally disordered regions in the future as the number of cancer studies increases, highlighting additional cancer types where novel drug design strategies targeting disordered regions are needed.

## Material and methods

### Identification driver regions in cancer-associated proteins

To collect mutation data, cancer mutations were retrieved from the v83 version of COSMIC[30] and the v6.0 version of TCGA. Mutations used from both databases included missense mutations, and in-frame insertions and deletions only. Mutations were filtered similarly to the procedure described in[31]. Mutations from samples with over 100 mutations were discarded to avoid the inclusion of hypermutated samples. Samples including a large number of mutations in pseudogenes or mutations indicated as possible sequencing/assembly errors in[67] were also discarded. Redundant samples were also filtered out. Mutations falling into positions of known common polymorphisms[68] were filtered. The final set of COSMIC mutations used as an input to region identification consists of 599,137 missense mutations, 4,189 insertions and 12,670 deletions from 253,568 samples. The final set of TCGA mutations used as an input to region identification consists of 274,109 missense mutations, 2,775 insertions and 2,900 deletions from 7,058 samples.

Driver regions were identified using iSiMPRe[31] with the filtered mutations from COSMIC and TCGA, separately. Then, regions obtained from COSMIC and TCGA mutations were merged, and p-values for significance were kept from the dataset with the higher significance. Only regions with high significance, with p-values lower than 10^−6^, were kept.

### Structural categorization of driver regions

Regions were assigned ordered or disordered status based on the structural annotation of the corresponding functional unit, incorporating experimental data as well as predictions. For this, we collected experimentally verified annotations for disorder from the DisProt[69] and IDEAL[70] databases, and for disordered binding regions from the DIBS[71] and MFIB[28] databases. We also mapped known PDB structures [72]). Structure of a monomeric single domain protein chain was taken as a direct evidence for order. In contrast, missing residues and mobile regions calculated for NMR ensembles using the CYRANGE method [73] were taken as indication of disorder. Pfam families annotated as of domain type were considered as ordered, while families annotated as disordered were assigned as disordered. All these types of evidence were extended by homology transfer.

Pfam entities with no instances overlapping with any protein regions with a clear structural designation, were annotated using predictions, together with protein residues not covered by known structural modules. Such protein regions were defined as ordered or disordered using predictions from IUPred[74,75] and ANCHOR[76,77]. Residues predicted to be disordered or to be part of a disordered binding region, together with their 10 residue flanking regions were considered to form disordered modules. Regions shorter than 10 residues were discarded. Regions annotated as disordered were also checked using additional prediction methods using the MobiDB database [78] and structure prediction using HHPred [79]. The final ordered/disordered status of the identified regions was based on manual assertion taking into account information from the literature, if available (Supplementary Table S1). For the disordered regions, the level of supporting information for the disordered region is also included (Supplementary Table S2).

### System-level analyses

Gene Ontology terms (GO[80,81]) were used to quantify interaction capabilities, involvement in various biological processes, molecular toolkits, and hallmarks of cancer. In each case a separate collection of GO terms (termed GO Slim) was compiled. Each GO Slim features a manual selection of GO terms that are independent from each other, meaning that they are neither child or parent terms of each other. Terms were assigned a level showing the fewest number of successive parent terms that include the root term of the ontology namespace (considered to be level 0).

GO term enrichments in a set of proteins were calculated by first obtaining expected values. Expected mean occurrence values for GO terms together with standard deviations were calculated by assessing randomly selected protein sets from the background (the full human proteome) 1,000 times. The enrichment in the studied set is expressed as the difference from the expected mean in standard deviation units.

GO for molecular toolkits: biological_process terms attached to proteins with identified regions were filtered for ancestry. The resulting set was manually filtered, yielding 93 terms, which were manually grouped into 16 toolkits. Enrichments for toolkits were calculated as the ratio of the sum of expected and observed values for individual terms. Individual terms and enrichments for each toolkit are shown in Supplementary Table S3.

GO Slim for assessing interaction capacity: terms from levels 1-4 from the molecular_function namespace were filtered for ancestry and only the more specific terms were kept. I.e. terms from levels 1-3 were only included if they have no child terms. Only terms describing interactions containing the keyword ‘binding’ were kept. Individual terms are shown in Supplementary Table S4.

GO for the assessment of process overlaps: terms from levels 1-4 from the biological_process namespace were filtered for ancestry and only the most specific terms were kept. Only those terms were considered that were attached to at least one protein from the set studied (full human proteome, ordered drivers, or disordered drivers). Individual terms are shown in Supplementary Table S5.

GO for hallmarks of cancer: Terms were chosen from the biological_process namespace via manual curation using the GO annotations of known cancer genes as a starting point. Terms were only kept if they showed a significant (p<0.01) enrichment on proteins in the full census cancer driver set compared to randomly selected human proteins. Individual terms and enrichments for each hallmark are shown in Supplementary Table S6.

To characterize the network properties of the selected examples, binary protein-protein interactions for the human proteome were downloaded from the IntAct database[82] on 06/05/2018. Data were filtered for human-human interactions, where interaction partners were identified by UniProt accessions. Interactions from spoke expansions were excluded. Interactions were kept in an undirected way. (Values for disordered drivers are quoted in Supplementary Table S2).

## Data availability

The authors declare that the data supporting the findings of this study are available within the paper and its supplementary information files.

## Acknowledgements

The authors thank Mark Adamsbaum and Drs Toby J. Gibson, Péter Tompa and László Buday for the critical reading of and their constructive comments on the manuscript.

## Funding

This work was supported by the “Lendület” grant from the Hungarian Academy of Sciences (LP2014-18) (Z.D.), OTKA grants (K108798 and K129164) (Z.D.) and the grant PD-120973 (A.Z) of the National Research, Development and Innovation office of Hungary and the EMBO|EuropaBio fellowship 7544 (B.M.).

## Competing Interests

The authors declare no competing interests.

## Supplementary material

### Supplementary figures

**Supplementary Figure S1.**
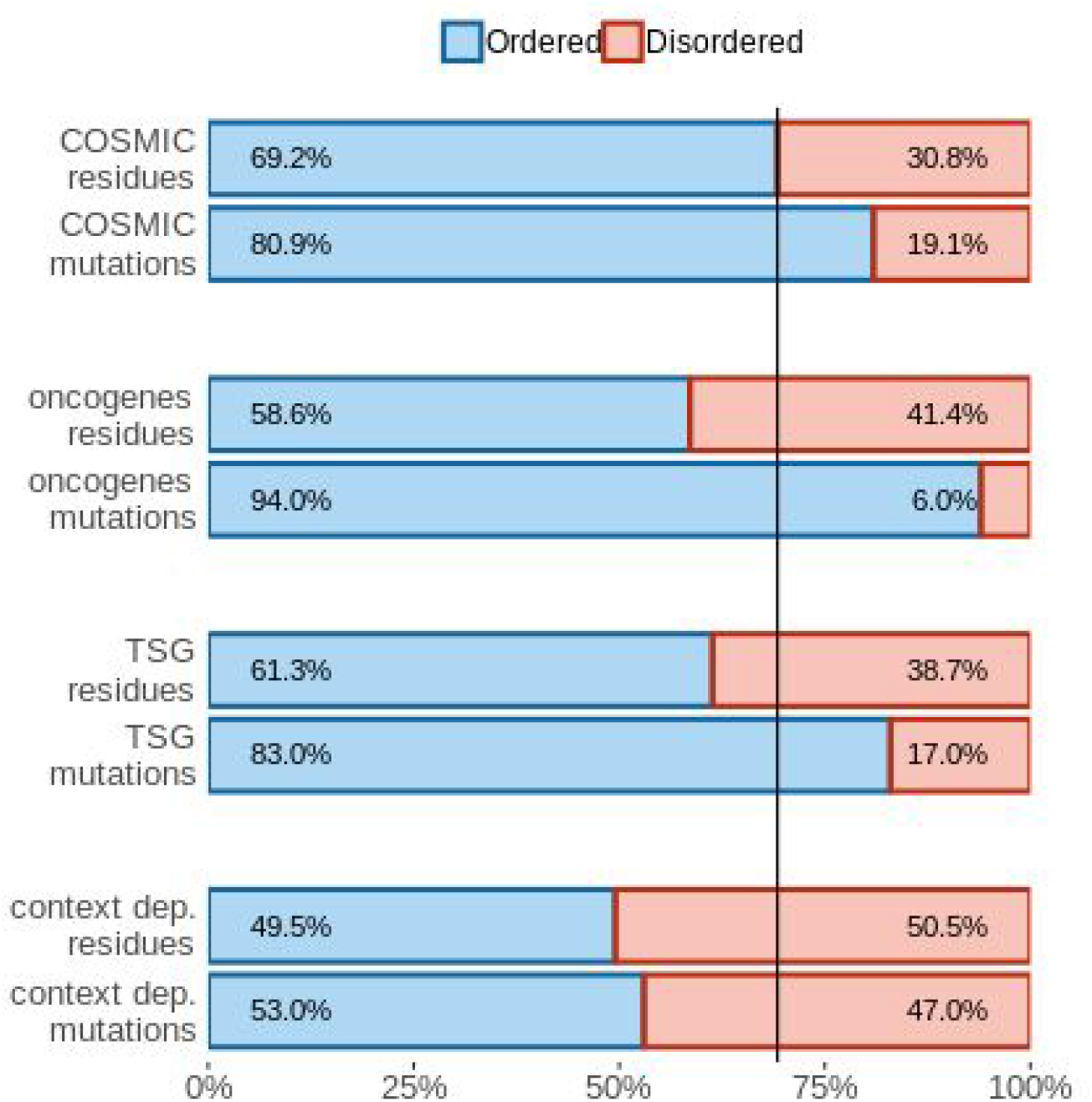
The distribution of residues and cancer mutations. The distribution of residues and COSMIC cancer mutations in the human proteome and various cancer driver classes. The vertical line marks the ratio of ordered and disordered residues in the human proteome, corresponding to the expected ratio of randomly occurring mutations.

**Supplementary Figure S2.**
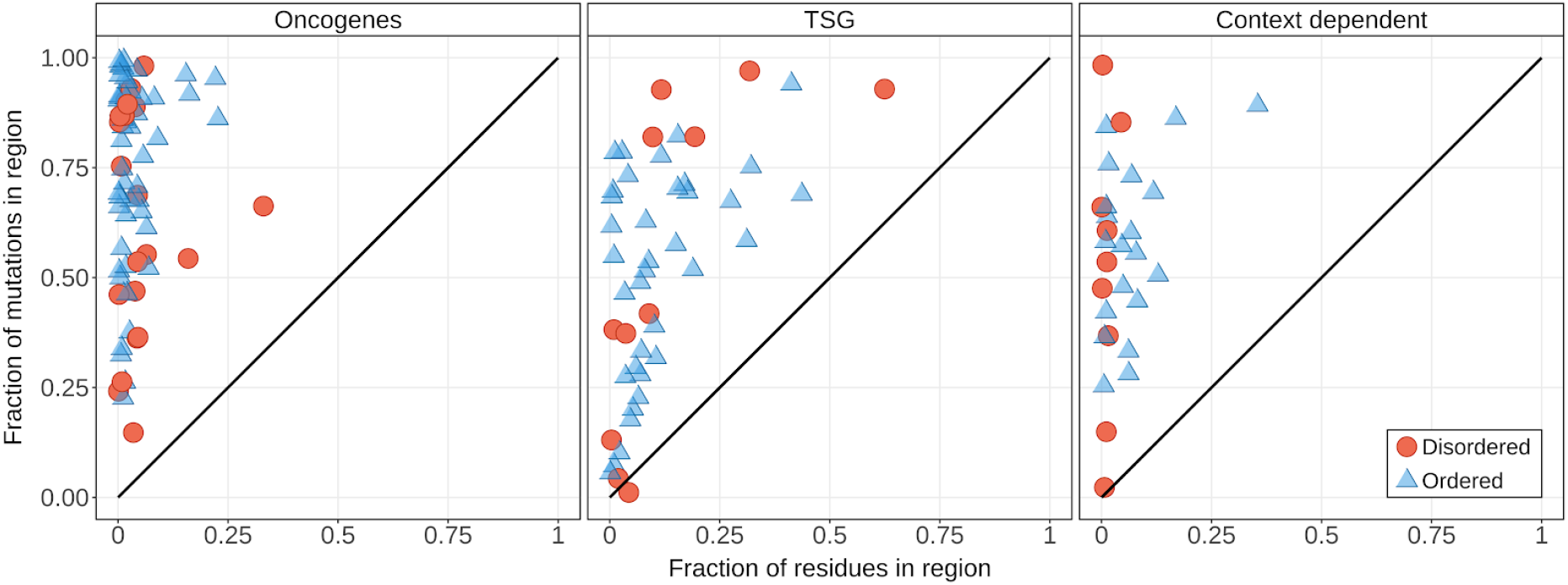
Identified regions are compact functional units. The fraction of mutations residing in the identified regions vs the fraction of residues in the regions for oncogenes, tumor suppressor genes and context-dependent genes. Regions in oncogenes encompass less than 10% of the sequence. Tumor suppressor genes (TSGs) are most often modulated via non-localized truncating mutations, affecting larger regions or the whole of the protein. However, the clustering of localized point mutations covering an extended region of the protein is a frequent hallmark of TSGs as well, making them largely identifiable using point mutations alone. In accord, several TSGs also harbor identified driver regions. While these regions are larger in size compared to that of oncogenes, they still cover only less than 20% of the sequence in most cases. In accordance with their dual nature, regions found inside context-dependent proteins lie between regions in oncogenes and TSGs in terms of length.

**Supplementary Figure S3.**
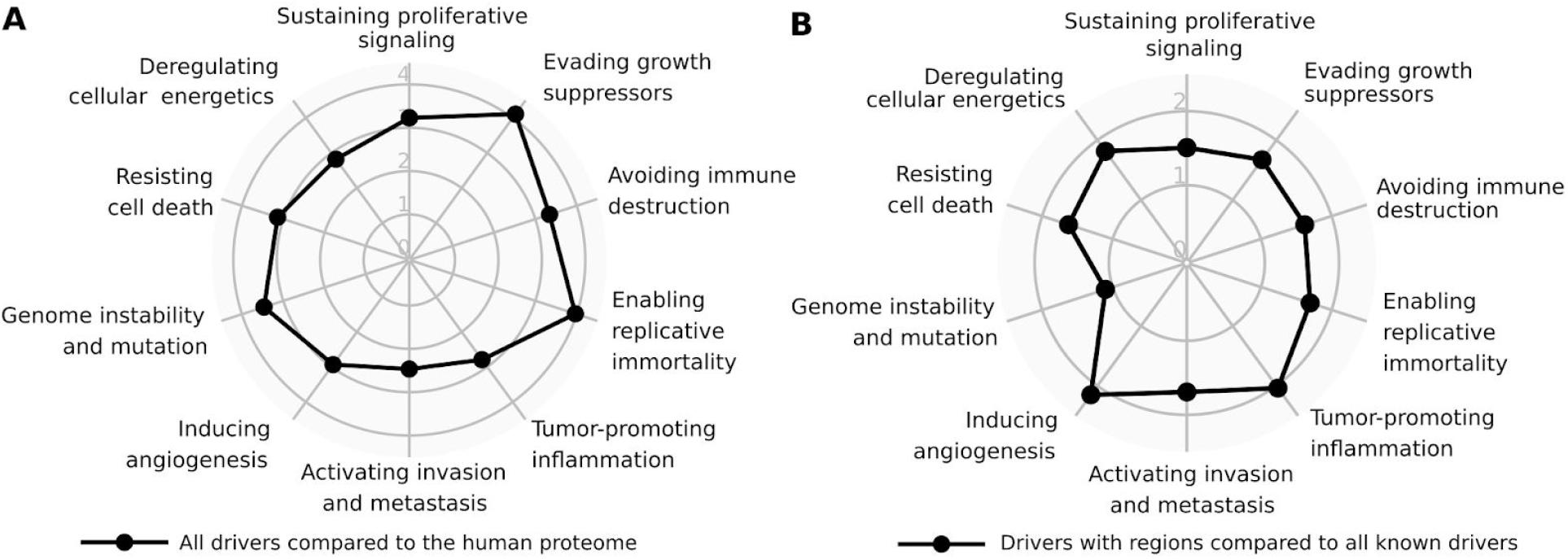
Overrepresentation of cancer hallmarks. A. Overrepresentation of hallmarks of cancer for all driver genes compared to the human proteome. B. Overrepresentation of hallmarks of cancer for driver genes with identified regions compared to all driver genes.

**Supplementary Figure S4:**
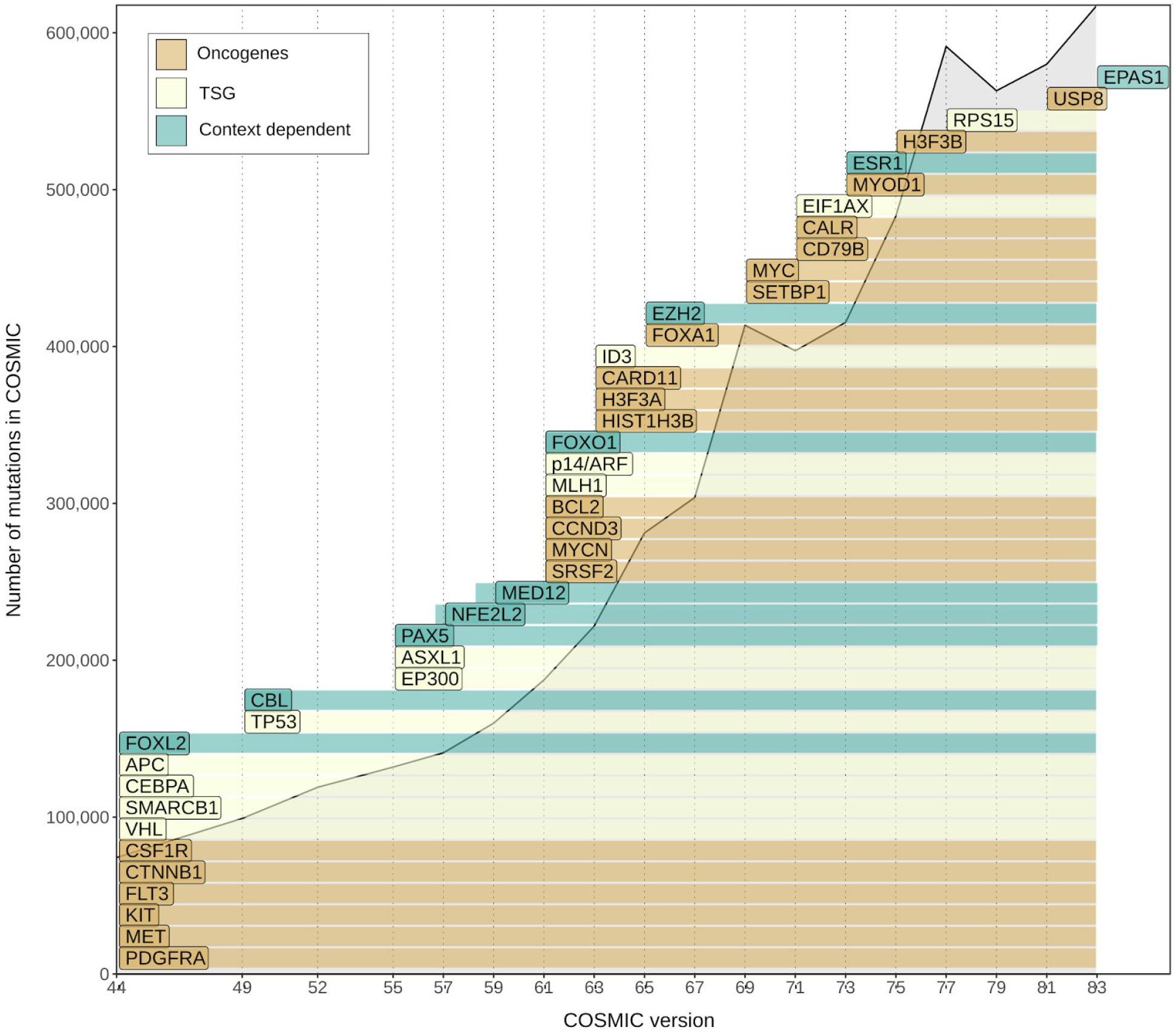
Growth of the list of disordered cancer drivers across various versions of the COSMIC database.

### Supplementary table legends

**Table S1**. List of regions identified using iSiMPRe, based on both COSMIC and TCGA mutations.

**Table S2**. Identified disordered driver genes with all annotations.

**Table S3**. Gene Ontology terms used in the quantification of molecular toolkits used by cancer driver genes.

**Table S4**. Gene Ontology terms used in the quantification of interaction capabilities.

**Table S5**. Gene Ontology terms used in the quantification of biological process overlaps.

**Table S6**. Gene Ontology terms used in the quantification of hallmarks of cancer.

## Notes

### Competing Interest Statement

The authors have declared no competing interest.

### Summary of Updates

Minor corrections

